# Genetic and brain similarity independently predict childhood anthropometrics and socioeconomic markers

**DOI:** 10.1101/2023.08.18.553837

**Authors:** Andreas Dahl, Espen M. Eilertsen, Sara F. Rodriguez-Cabello, Linn B. Norbom, Anneli D. Tandberg, Esten Leonardsen, Sang Hong Lee, Eivind Ystrom, Christian K. Tamnes, Dag Alnæs, Lars T. Westlye

**Affiliations:** Department of Psychology, University of Oslo, Oslo, Norway; NORMENT, Division of Mental Health and Addiction, Oslo University Hospital & Institute of Clinical Medicine, University of Oslo, Oslo, Norway; Research Center for Developmental Processes and Gradients in Mental Health (PROMENTA), Department of Psychology, University of Oslo, Oslo, Norway; Department of Psychiatric Research, Diakonhjemmet Hospital, Oslo, Norway; Australian Centre for Precision Health, UniSA Allied Health & Human Performance, University of South Australia, Adelaide, Australia; South Australian Health and Medical Research Institute (SAHMRI), University of South Australia, Adelaide, Australia; Department of Mental Disorders, Norwegian Institute of Public Health, Oslo, Norway; KG Jebsen Center for Neurodevelopmental Disorders, University of Oslo, Norway

## Abstract

Linking the developing brain with individual differences in clinical and demographic traits is challenging due to the substantial interindividual heterogeneity of brain anatomy and organization. Here we employ a novel approach that parses individual differences in both cortical thickness and common genetic variants, and assess their effects on a wide set of childhood traits. The approach uses a linear mixed model framework to obtain the unique effects of each type of similarity, as well as their covariance, with the assumption that similarity in cortical thickness may in part be driven by similarity in genetic variants. We employ this approach in a sample of 7760 unrelated children in the ABCD cohort baseline sample (mean age 9.9, 46.8% female). In general, significant associations between cortical thickness similarity and traits were limited to anthropometrics such as height (r^2^ = 0.11, SE = 0.01), weight (r^2^ = 0.12, SE = 0.01), and birth weight (r^2^ = 0.19, SE = 0.01), as well as markers of socioeconomic status such as local area deprivation (r^2^ = 0.06, SE = 0.01). Analyses of the contribution from common genetic variants to traits revealed contributions across included outcomes, albeit somewhat lower than previous reports, possibly due to the young age of the sample. No significant covariance of the effects of genetic and cortical thickness similarity was found. The present findings highlight the connection between anthropometrics as well as socioeconomic factors and the developing brain, which appear to be independent from individual differences in common genetic variants in this population-based sample. The approach provides a promising framework for analyses of neuroimaging genetics cohorts, which can be further expanded by including imaging derived phenotypes beyond cortical thickness.

## Introduction

Mapping individual differences in brain morphology and their associations with relevant clinical and demographic traits has been described as one of the fundamental challenges of neuroscience (Giedd and Rapoport 2010; Lashley 1947). This task is particularly challenging in young individuals, as the structure of the brain changes rapidly when they progress through different stages of development (Mills et al. 2021). A morphological measure that has received extensive attention is the thickness of the cortex, both due to its potential sensitivity to age (Frangou et al. 2022) and clinical conditions (Hettwer et al. 2022). Cortical thickness can be estimated from magnetic resonance imaging (MRI) data with reasonable accuracy (Fuhrmann et al. 2022). However, reported associations between apparent cortical thickness and observable traits in children and adolescents have largely been inconclusive (Marek et al. 2022). This lack of robustness may in part be attributed to a methodological reliance on average effects that inadequately accounts for the individual heterogeneity of cortical structure and development, instigating a call for new approaches that better capture this variability (Foulkes and Blakemore 2018; Westlin et al. 2023).

In the field of genetics, leveraging the inherent genetic similarity among individuals to explore the relationship between their genetic make-up and observable traits has revealed novel insight into the associations between genetic factors and human traits (Tam et al. 2019). For this, a genomic relatedness matrix (GRM) can be constructed (J. Yu et al. 2006) by estimating the pairwise resemblance of individuals in a sample based on genome-wide single nucleotide polymorphisms (SNPs; Yang et al. 2010). This GRM can then be integrated into linear mixed models (LMMs) along with phenotypic traits, enabling the estimation of the proportion of phenotypic variance attributed to genetics (commonly known as SNP-based heritability). This approach, often referred to as genome-based restricted maximum likelihood (GREML), has been successfully applied to large cohorts of young individuals to estimate SNP-based heritability for complex behavioral traits, such as academic performance, psychological distress, and externalizing behavior (Cheesman et al. 2020; Donati et al. 2021; Eilertsen et al. 2022; Jami et al. 2022). The application of GREML in neuroscience has been limited compared to other fields (Trzaskowski et al. 2014), and between-subject variability is commonly considered as error. There is a growing recognition of the potential benefits of incorporating individual variability caused by genetics to enhance our understanding of the relationship between the developing cortex and observable traits (Z. Yu et al. 2022).

Applying a similar approach to GREML, Sabuncu et al. (2016) reported that the phenotypic variance of both clinical (e.g., diagnosis of mental illness) and non-clinical traits (e.g., cognition) could be significantly explained by whole-brain morphology, specifically a composite of multiple gray- and white matter measures in adults. This approach has been termed *morphometricity (Sabuncu et al. 2016)*, or *trait morphometricity* (Fürtjes et al.(2023). In simplified terms, this approach entails estimating one or more measures of brain morphology, such as cortical thickness and cortical surface area. Next, the pairwise resemblance across all vertices or regions of interest (ROIs) of one or more such morphological measures is calculated across all individuals, resulting in a brain-morphological similarity matrix, analogous to a GRM. The resulting matrix is then used in LMMs, yielding an estimate of the proportion of phenotypic variance attributed to brain morphology. This approach has been expanded by Couvy-Duchesne et al. (2020) and Fürtjes et al. (2023), showing that similarity matrices based on morphological measures explain significant proportions of variance across different groups of traits, such as anthropometrics (e.g. BMI), cognition, markers of socioeconomic status (SES) and health behaviors (Couvy-Duchesne et al. 2020). Importantly, comparisons have shown that similarity-based approaches consistently outperformed conventional univariate association analyses, both in terms of power to detect effects and in explained trait variability, in clinical and non-clinical traits (Sabuncu et al. 2016).

The current study expands on this work by investigating both the morphometricity and SNP-based heritability of a wide array of traits in a large sample of US children from the ABCD cohort baseline sample. Our novel contribution will be threefold. First, we are not aware of any previous study investigating the morphometricity of traits in younger individuals. It is conceivable that this approach might manifest differently in children compared to adults. We will restrict our approach to cortical thickness, which shows marked changes during development (Fuhrmann et al. 2022), and is reasonably robust against confounds such as head size and total brain volume (Barnes et al. 2010). Second, by assessing both morphometricity and SNP-based heritability within the same LMM framework, we estimate the observed trait variance that can be explained by both genomic and morphological effects, i.e. a combined genome-morphometric analysis. Our approach also allows for the exploration of the covariance between the genomic and morphological effects, using the CORE GREML approach (Zhou et al., 2020). The purpose of this is twofold: Traditionally, REML estimation assumes independence between random effects. However, cortical morphology has been shown to be heritable in both adults and younger individuals (Fernandez-Cabello et al. 2022; Shadrin et al. 2021; van der Meer et al. 2020), potentially biasing estimates. By utilizing the CORE GREML approach, we can account for potential dependencies between genomic and morphological random effects, resulting in a more accurate estimation of their respective contributions. In addition, the inclusion of a third term describing the covariance allows for the delineation of the unique contributions of genomic and morphological effects on the trait of interest. This allows us to assess if the potential covariance of their effects manifest differently depending on the trait under investigation

## Materials and Methods

### Participants

The full sample for the main analysis following MRI and genetics quality control (QC; see below) consisted of data from 7760 individuals (mean age 9.9 years, 46.8% females) obtained from the Adolescent Brain Cognitive Development Study (ABCD) annual data release 3.0 (http://dx.doi.org/10.15154/1523041). The ABCD study (https://abcdstudy.org/) is an ongoing longitudinal developmental study (Volkow et al. 2018) following participants from age ∼10 to age ∼20, with bi-annual collection of neuroimaging data. Only data from the baseline session was included in the current analyses.

### Ethical approval

The review and approval of the ABCD research protocol was handled by a central Institutional Review Board at the University of California, San Diego (Auchter et al. 2018). Informed consent was given by parents or guardians and assent was given by children before participation. The present project is registered in the NIMH Data Archive as project number 1467 (doi: 10.15154/1524691), available for registered and authorized users (Request #7474, PI: Westlye). The current project has also been approved by the Norwegian Regional Committee for Medical and Health Research Ethics (REC; #2019/943).

### Genetic data -genomic relatedness matrix

Genotyped data was provided by the ABCD consortium, specifically from the Genomics sample_03 (https://nda.nih.gov/study.html?id=1299). A full description of the collection and handling of genotyped data can be found at https://nda.nih.gov/experimentView.html?experimentId=1194. QC was performed by the ABCD consortium using the RICOPILI pipeline (Lam et al. 2020). Robust relatedness estimates were generated from genotyped SNPs using the *pcrelate* function from GENESIS version 2.24.0 (10.18129/B9.bioc.GENESIS; Conomos et al. 2016), and converted into a GRM using the *pcrelateToMatrix* function from the same package. A GRM describes an estimate of the additive genetic relationship between individuals, where each off-diagonal entry denotes the estimated relatedness for a pair of individuals. It can be expressed as

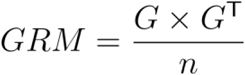

Where *GRM* is the resulting genomic relatedness matrix, *G* are columns of allele counts standardized to have a mean zero and a standard deviation of one, and is the number of SNPs. Before the final analysis, for pairs with a familial or cryptic relatedness of 0.05 and above, one individual was removed using the *grm-cutoff* function from GCTA version 1.93.0 (Yang et al. 2011), leaving the maximum possible sample size of non-related individuals.

### MRI QC and processing

A full description of ABCD MRI collection and acquisition parameters is given in Casey et al. (2018). Participants that did not pass the recommended image inclusion criteria provided by the ABCD consortium were removed from the sample (imgincl_t1w_include == 0; see http://dx.doi.org/10.15154/1523041 for full details of the QC procedure). T1-weighted MRI data from participants that passed the QC were processed using FreeSurfer 7.1 (surfer.nmr.mgh.harvard.edu). Cortical thickness was computed vertex-wise, as coarser atlas-based ROIs may carry insufficient spatial information for reliable estimates of the morphometricity of traits (Fürtjes et al. 2023). Individual cortical thickness surfaces were registered to a common template (fsaverage) and smoothed using a 15 mm full width at half maximum (FWHM) gaussian kernel. Non-cortical vertices belonging to the medial wall were excluded, leaving a total of 299 879 vertices across both hemispheres for each participant.

To account for scanner-related confounds, a ComBat harmonization procedure was implemented in neuroCombat version 1.0.13 in R (https://github.com/Jfortin1/neuroCombat_Rpackage), using an empirical Bayes location-shift model for all 28 scanners (see supplementary figure 1). All outcome measures were added as covariates for the harmonization procedure to preserve the presumed biological variability of trait outcomes. The resulting harmonized cortical thickness measures are a linear combination of the variables of interest and a scanner-specific residuals modulated by both additive and multiplicative scaling factors (Fortin et al. 2018).

**Figure 1.**
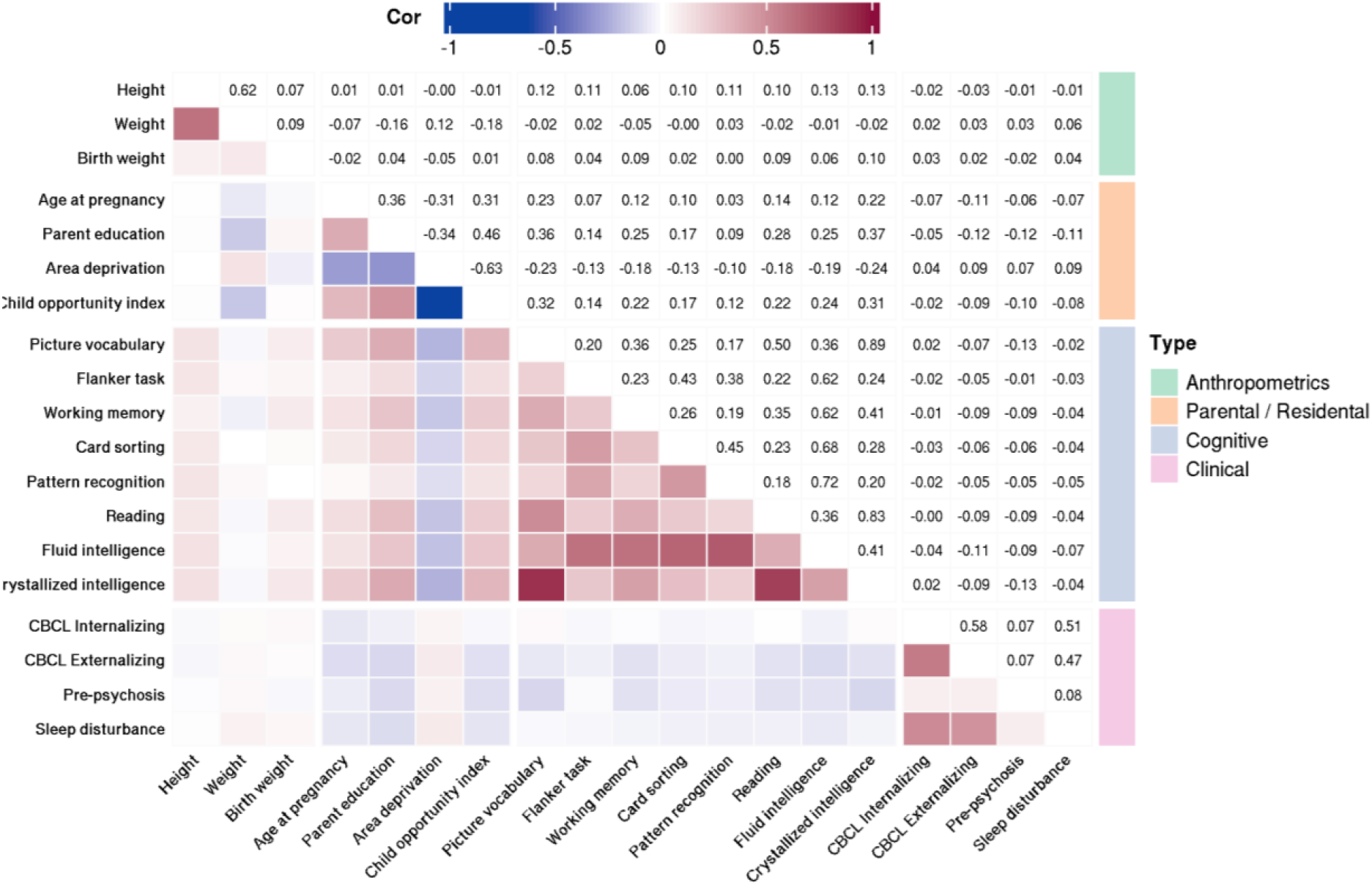
Correlations of all included outcome variables. The upper and lower triangular represent the same values, numerically (upper) and color coded (lower).

### Brain similarity

To determine morphological similarity based on cortical thickness, we calculated the cross-product of the transpose of a matrix containing all vertices of all participants. The formula is equivalent to the calculation of GRM, i.e.

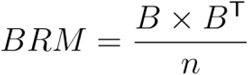

Where *BRM* is the resulting *brain relatedness matrix* (BRM), with each off-diagonal element describing the degree of similarity in morphology between two individuals, *B* is a matrix.containing centered and scaled measures of cortical thickness for all vertices, standardized to have mean zero and standard deviation of one and *n* is the total number of voxels.

### Covariance between the effects of genomic relatedness and brain relatedness

To investigate the covariance of the effects of brain measures and genomic data, we used the CORE GREML approach developed by Zhou et al. (2020). CORE GREML extends the concept of genome-based restricted maximum likelihood (GREML) by enabling the estimation of the covariance between two random effects through the product of the Cholesky decomposition of the two relatedness matrices. The detailed procedure can be found in Zhou et al. (2020). Briefly, the GRM and BRM matrices were transformed to be positive-definite and subjected to Cholesky decomposition. Subsequently, the product matrix of the Cholesky decompositions of the GRM and BRM was calculated. All the necessary steps of this procedure were implemented in MTG2 version 2.22 (Lee and van der Werf 2016). Estimates of model parameters for the covariance were obtained by fitting the product matrix, along with the GRM and the BRM, in an LMM (see Model 2 below).

### Outcome measures

All outcome measures were taken from ABCD data release 3.0. and handled in R version 4.0.0 (https://cran.r-project.org). We included outcome measures from four different domains: anthropometric, parental / residential, cognitive, and clinical (e.g. potential early markers of mental illness). Detailed descriptions of included instruments are given in Table 1. Pearson correlations of all included outcomes are given in Figure 2. Anthropometrics such as height and weight are highly heritable (Momin et al. 2023), and previously shown considerable levels of morphometricity in adults, with cortical morphology accounting for approximately 20% of the variation in body mass index (Fürtjes et al. 2023). However, heritability estimates of anthropometric measures tend to be lower during childhood and adolescence (Jelenkovic et al. 2016). It remains uncertain if estimates of morphometricity would be equally reduced. As a growing body of evidence demonstrates associations between perinatal and early-life factors and later brain development (Alnæs et al. 2020; Walhovd et al. 2023), we also included weight at birth.

**Table 1.**
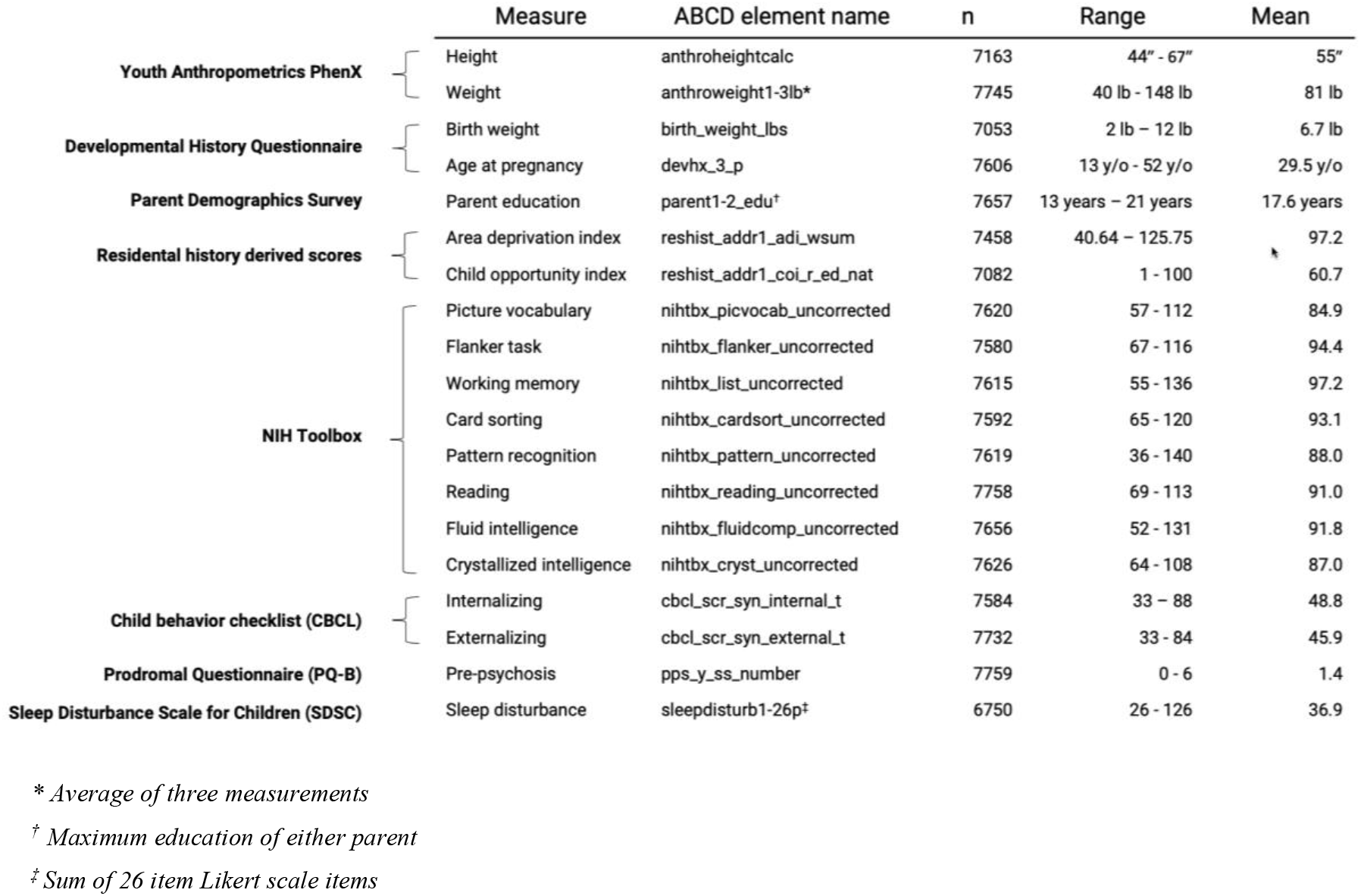
Included outcome variables. n equals the final number of available data following genetic and MRI QC and outlier removal.

**Fig. 2.**
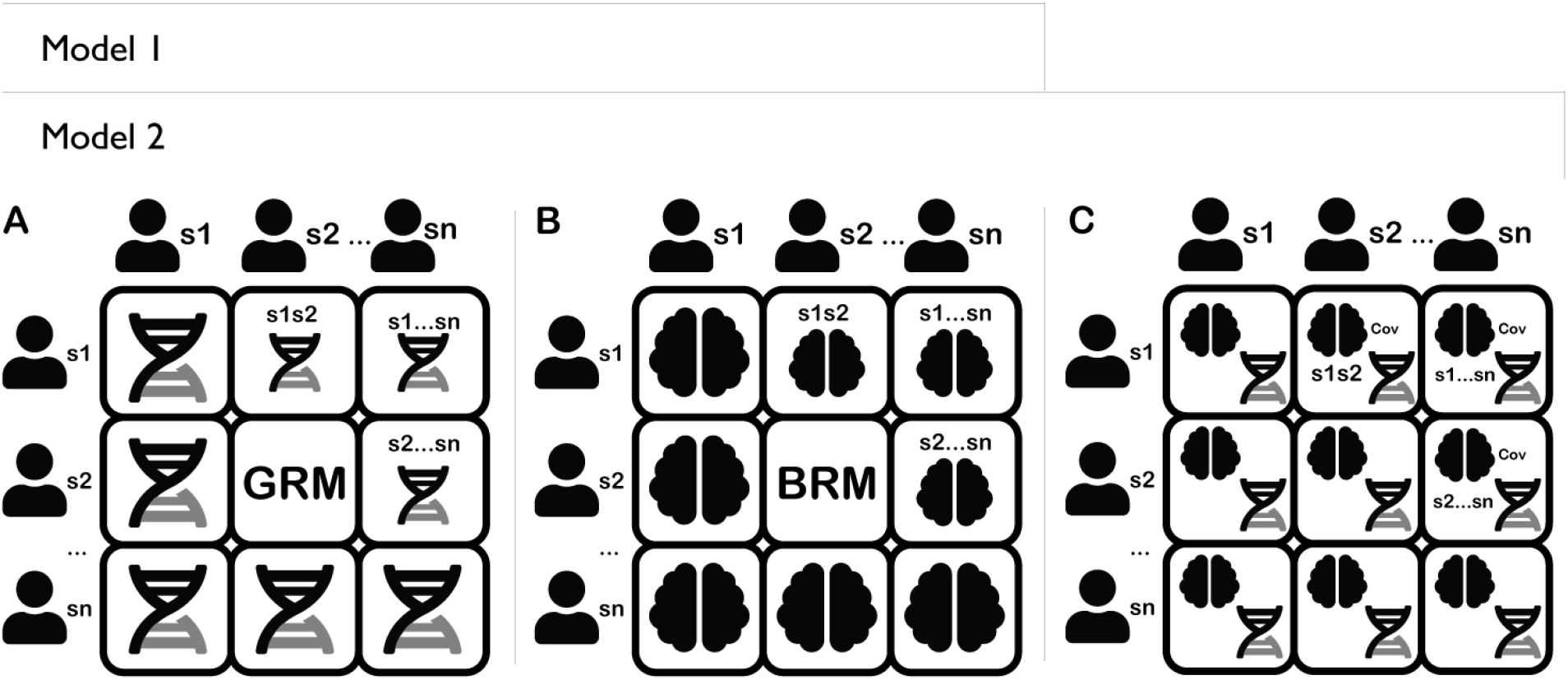
Illustration of random effects included in Model 1 and Model 2. A: Genomic relatedness matrix. B: Brain relatedness matrix. C: Covariance of effects of A and B.

Measures of cognition and general intelligence were included due to their clinical and functional relevance and links to both cortical development and genetics (Estrada et al. 2019). For the remaining included measures, we attempted to capture the associations between morphology, genetic influences, and the family and local environment. This includes markers of socioeconomic status, which have previously been associated with brain imaging derived phenotypes in the ABCD sample (Alnæs et al. 2020). Lastly, we included measures of early signs of mental illness, including externalizing and internalizing symptoms.

Outcome scores more than four median absolute deviations from the median were set to missing (Leys et al. 2013). Following this, histograms of outcome distributions were inspected manually, resulting in four weight measurements, all below 40lbs/18kg., being set to missing. For each outcome variable missing data was removed before being ordered-quantile-normalized using the *bestNormalize* package version 1.8.2. in R (https://cran.r-project.org/package=bestNormalize).

### Data analysis

First, we calculated the overall Pearson’s correlation between the off-diagonal elements of the GRM and the BRM. This correlation provided insights into the similarity or dissimilarity between the two matrices, irrespective of their associations with specific traits. Due to the extensive number of elements, we report descriptive statistics only.

Second, morphometricity and SNP-based heritability estimates were obtained using two separate restricted likelihood random-effects (REML) models for each of the 19 phenotypes (Fig 2.). The first model is

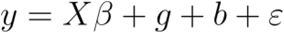

where *y* is the trait of interest, *X* is an incident matrix for the fixed effects *β*; age, sex, genotype batch, and the first 20 principal components of the GRM and four genetic ancestry factors (GAFs) to account for population stratification and ancestry (East Asian, African, American, European).*g* is the random genomic effects and *b* is the random effects of morphological measures. Then, the variance and covariance *y* of can be written as

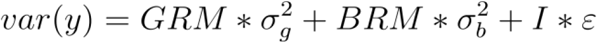

The second model (i.e. CORE GREML), denoted as Model 2, is the same as in Model 1 except for the addition of the covariance term in the variance and covariance of *y*, which can be written as

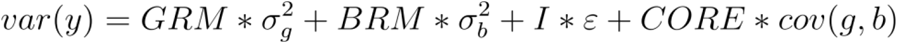

where *CORE* is the product of the Cholesky decomposition of the GRM and the BRM, and *cov*(*g,b*) is the covariance between genomic and morphological effects.

The final calculation of morphometricity (*m*^2^) is equivalent to conventional heritability (*h*^2^) calculation, e.g.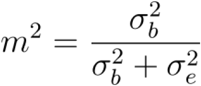 is the same as 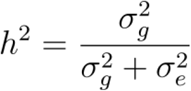

Estimates of standard error of *m*^2^ and *h*^2^ were obtained using the Delta method (Oehlert 1992)). Reported p-values are based on Wald tests with one degree of freedom under the null hypothesis that the variance component is zero, implemented in MTG2. Likelihood ratio tests with 1 degree of freedom were performed to determine if the addition of the covariance term significantly improved model fit for a given trait (CORE GREML). This was done only on traits where both the GRM and BRM contributed significantly to trait variance. It gives that if the contribution of two random effects to variation in a trait is negligible, their covariance would also be negligible. The correlation estimates reported in Table 2 is the correlation of the two random effects *g* and *b*, obtained as in Zhou et al. (2020) by scaling the covariance by the square root of the product of the variance of the two random effects, i.e.

**Table 2:** Outcome of log likelihood comparisons of Model 1 and Model 2 and correlations of random effects.

| Trait | $h^2$ CORE GREML | | | $m^2$ CORE GREML | | | Log likelihood | | | Correlation of random effects | | |
| --- | --- | --- | --- | --- | --- | --- | --- | --- | --- | --- | --- | --- |
|  | Estimate | SE | p-value | Estimate | SE | p-value | w/o CORE GREML | CORE GREML | p-value diff. (df=1) | Corr. | SE | p-value |
| Height | 0.188 | 0.045 | <0.001* | 0.110 | 0.014 | <0.001* | -2710.69 | -2710.69 | 0.968 | 0.005 | 0.132 | 0.968 |
| Weight | 0.134 | 0.043 | 0.005* | 0.116 | 0.014 | <0.001* | -3057.03 | -3056.38 | 0.254 | 0.171 | 0.154 | 0.956 |
| Parent education | 0.138 | 0.049 | 0.009* | 0.026 | 0.008 | 0.002* | -2329.61 | -2328.44 | 0.125 | 0.407 | 0.292 | 0.956 |
| Area deprivation | 0.145 | 0.049 | 0.007* | 0.059 | 0.011 | <0.001* | -2743.15 | -2742.56 | 0.276 | 0.203 | 0.197 | 0.956 |
| Child opportunity index | 0.163 | 0.050 | 0.004* | 0.020 | 0.007 | 0.009* | -2150.07 | -2149.77 | 0.445 | 0.206 | 0.281 | 0.956 |
| Picture vocabulary | 0.191 | 0.050 | <0.001* | 0.016 | 0.006 | 0.013* | -2749.55 | -2749.19 | 0.395 | -0.226 | 0.260 | 0.956 |
| Working memory | 0.120 | 0.049 | 0.022* | 0.025 | 0.007 | 0.002* | -3287.35 | -3287.33 | 0.827 | -0.063 | 0.286 | 0.956 |
| Pattern recognition | 0.119 | 0.049 | 0.022* | 0.018 | 0.006 | 0.009* | -3529.75 | -3529.62 | 0.604 | -0.182 | 0.329 | 0.956 |
| Reading | 0.205 | 0.050 | <0.001* | 0.017 | 0.006 | 0.009* | -3206.03 | -3205.93 | 0.663 | -0.109 | 0.247 | 0.956 |
| Fluid intelligence | 0.180 | 0.048 | <0.001* | 0.033 | 0.008 | <0.001* | -3091.11 | -3091.09 | 0.857 | 0.038 | 0.211 | 0.956 |
| Crystallized intelligence | 0.237 | 0.050 | <0.001* | 0.022 | 0.007 | 0.003* | -2745.79 | -2745.56 | 0.496 | -0.143 | 0.207 | 0.956 |

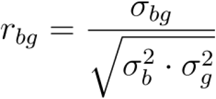

The sampling variance of the correlation estimates was obtained using the Delta method with variance and covariance terms from Model 2 (CORE GREML).

All reported p-values for variance components, likelihood ratio tests and correlations were adjusted for multiple tests by using false discovery rate (FDR; Benjamini and Hochberg 1995).

## Results

### Gross association of GRM and BRM elements

Correlation analyses revealed a near-zero association between the off-diagonal elements of the GRM and the BRM (r = 0.0015; 95% confidence interval [CI] = 0.0012, 0.0019), indicating that similarity in cortical morphology in children is not associated with genomic similarity (Figure 3).

**Fig. 3.**
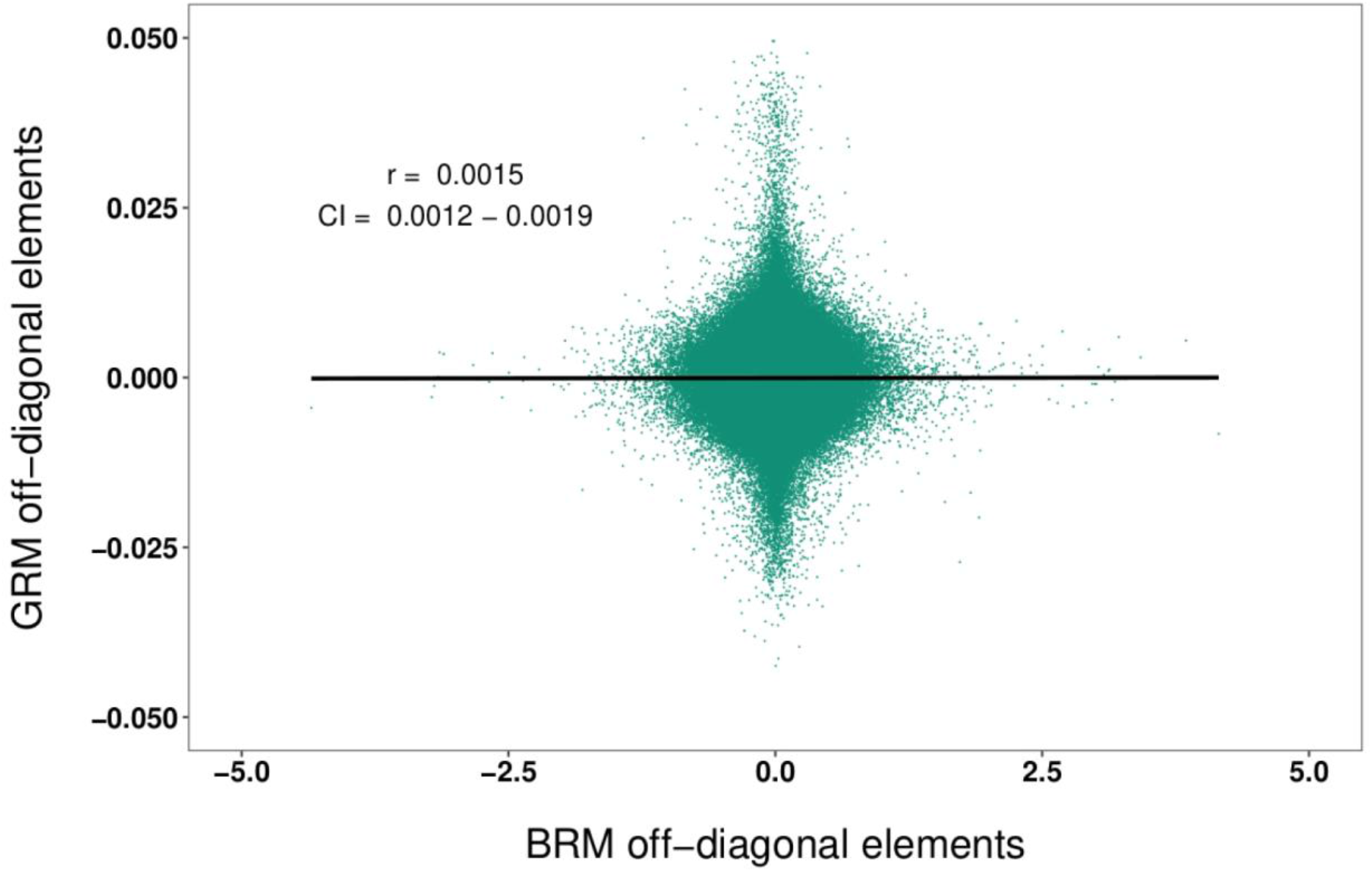
Pearson correlation of the off-diagonal elements of the GRM and the GRM. Scatter shows a random selection of 1 000 000 associations, values indicate the overall r and CI for all 30 104 920 associations.

#### Model 1

The full results of Model 1 analyses are presented in Fig. 4 and Supplementary Table 1. The estimates of SNP-based *h*^2^ were significantly different from zero for the majority of included traits. However, it should be noted that the contribution of genetic factors was generally modest and estimated SNP-based *h*^2^ did not exceed 0.30 for any trait. The highest estimates were found for the NIH Toolbox crystallized intelligence composite score (*h*^2^ = 0.23), the NIH Toolbox reading task (*h*^2^ = 0.20), and height (*h*^2^ = 0.19). Genomic similarity was not significantly associated with birth weight, mother’s age at pregnancy, the NIH Toolbox flanker task, internalizing and externalizing symptoms or sleep disturbance (all *h*^2^ < 0.1).

**Fig. 4:**
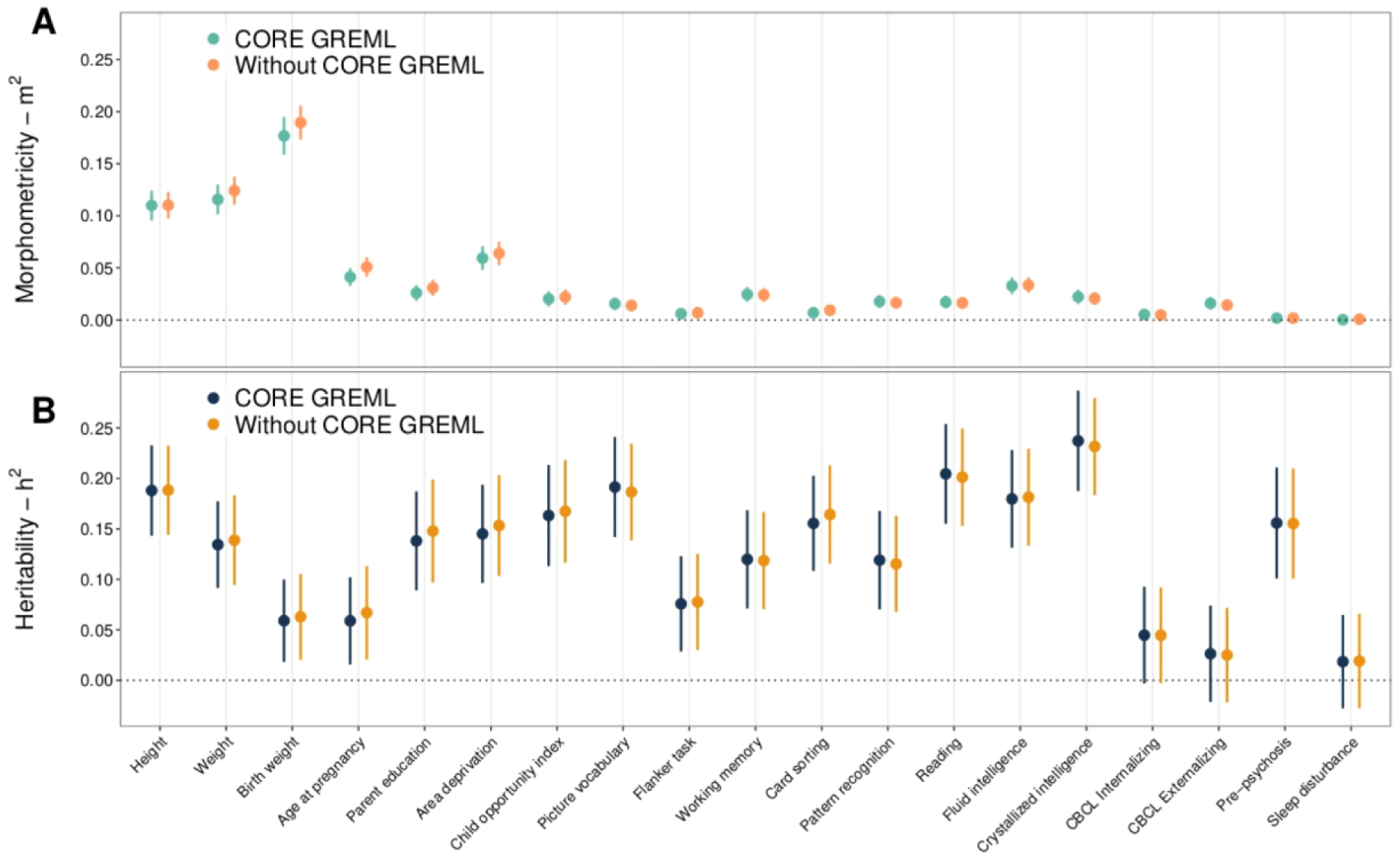
Outcomes of main analysis. (a) m^2^ estimates and SE for all included traits, either with or without the covariance (CORE GREML)-term included in the LMM. (b) h2 estimates and SE for all included traits, either with or without the covariance term included in the LMM.

The morphometricity analyses revealed associations between morphology and multiple traits of interest. Among these traits, the highest estimates of morphometricity (*m*^2^) were found for anthropomorphic traits, including birth weight (*m*^2^ = 0.19), current weight (*m*^2^ = 0.12) and height (*m*^2^ = 0.11). Significant *m*^2^ was also found for ten other traits of interest. However, it is important to note that the effects of these estimates were marginal, all below 5%, except for the area deprivation index (*m*^2^ = 0.06), and mothers’ age at pregnancy (*m*^2^ = 0.05).

#### Model 2

As evidenced by Figure 4, estimates of *m*^2^ and *h*^2^ did not show noticeable changes whether using GREML or CORE GREML. To further assess the significance of covariance, likelihood ratio tests were conducted with one degree of freedom for the ten traits where both genomic and morphological effects contributed significantly to variation in traits of interest (Table 2). The likelihood-ratio tests indicated that the addition of a third component describing the covariance between the genomic and morphological effects did not result in a significant change in the goodness of fit for any of the traits.

## Discussion

Investigations of associations between observable traits and brain structure among rapidly maturing children and adolescents often yield inconsistent results. In the present paper, we adopted methods from genetics and assessed the proportion of observable trait variance in children that can be explained by both similarity in common genetic variants and similarity in cortical thickness. Most included traits showed moderate heritability. However, beyond anthropomorphic traits, our analyses revealed generally weak associations between cortical thickness and included traits. Further, a novel approach to assessing the covariance between genomic and morphological effects revealed no strong interdependence, suggesting that their contributions were unique.

### Morphometricity

Our findings indicate the contribution of similarity in cortical morphology to the included traits were generally limited. This adds to recent literature suggesting that interindividual differences in cortical morphology share limited associations with behavioral differences among populations of normally developing children and adolescents (Genon, Eickhoff, and Kharabian 2022). This may also extend to adults, where previous estimates of strong brain-behavior associations from small-scale studies have proved difficult to replicate in large-scale population-based samples (Botvinik-Nezer and Wager 2022). Overall, our findings indicate that morphometricity is, as previously shown (Couvy-Duchesne et al. 2020; Fürtjes et al. 2023), reasonable for traits that are anthropometric in nature, such as height and weight, but this does not extend to psychopathology or cognitive functions in this young population based sample. Couvy-Duchesne et al. (2020) specifically probed the association between anthropometrics and morphometricity and found that the morphometricity of traits from multiple different categories, such as symptoms of mental disorders, were in part attributable to body size. Birth weight, however, appeared in our sample to be unrelated to current height or weight. This indicates that birth weight may have associations with cortical morphology that are independent of later body size. Although we cannot determine the directionality of effects in the present study, previous studies indicate that low birth weight is associated with an enduring pattern of accelerated brain maturation (Alnæs et al. 2020; Knickmeyer et al. 2017). In another recent study, Gilmore et al. (2020) showed that later heterogeneity of cortical thickness is largely present at 1 year of age, highlighting the lasting importance of neonatal characteristics on later brain development.

The finding that neighborhood socioeconomic conditions (as measured by the ADI) show significant, albeit somewhat limited, associations with cortical thickness supports previous research demonstrating that socioeconomic status is recognized in the child brain (Alnæs et al. 2020; Norbom et al. 2023). This association appears to go beyond population stratification and genetic ancestry, which were included as fixed effects in our models. The cause of this association is not known, but recent papers based on material from the ABCD study suggest that the association between SES and brain morphology is partly mediated by a lack of supportive stimulation and a lack of healthy food options more frequently found in lower compared to higher SES strata (Dennis, Manza, and Volkow 2022; Tomasi and Volkow 2021). We also observed that interindividual differences in cortical thickness was associated with maternal age at pregnancy. While our analysis does not inform us about the directionality of this effect, lower maternal age has previously been linked to lower SES strata (Moore et al. 1993; Restrepo-Méndez et al. 2015). It is possible that this association is confounded by birth weight, which showed moderate association with cortical morphology in our sample, and has previously been linked to both SES and maternal age (Restrepo-Méndez et al. 2015). However, in the present sample, the correlations between birth weight and SES markers were virtually non-existent, indicating that SES may have links to cortical morphology beyond gestational factors.

### Heritability analyses

We found that the majority of traits included were moderately heritable, which can be used as a reference for future investigations of SNP-based heritability in the ABCD study. However, we would like to acknowledge that some estimates are at the lower end compared to what is commonly reported. This is particularly true for height, with our estimate being approximately one third of what is typically found in adult populations (Yengo et al. 2022). The comparatively lower estimates of heritability may possibly be attributed to the age of the sample, as the heritability of many traits tends to be lower during childhood before increasing throughout adolescence (Bergen, Gardner, and Kendler 2007). This is also the case for height, with heritability estimates increasing dramatically from 11-12 years onwards (Jelenkovic et al. 2016). Another possible issue explaining the somewhat lower estimates of SNP-based heritability is the transethnic nature of the ABCD sample. Significant heterogeneity in either the genotype or the trait across different ethnic populations may cause a deflation of global SNP-heritability. This effect may be present even as population stratification and ancestry scores are added as covariates in LMMs (Li and Keating 2014). However, due to sample size constraints we did not consider it possible to run separate analyses for different ethnic ancestries.

### Covariance

We show that the covariance between the genomic and morphological effects on the trait of interest is not significantly different from zero, and these effects are largely independent. This finding has two implications. Firstly, it indicates that our estimates of heritability and morphometricity may not be affected by the covariance between these factors. Secondly, it suggests that while cortical morphology has been shown to be highly heritable (van der Meer and Kaufmann 2022), this does not necessarily translate into similar cortical thickness between individuals who share genomic similarity, at least not conditioned on the traits included in the present study. Conceivably, the covariance of the effects of genomic and cortical thickness similarity might present itself with age, as the influence of genetic factors on cortical morphology becomes stronger throughout adolescence (Schmitt et al. 2014). However, the lack of relationship between SNP-based and cortical thickness-based similarity, as evidenced by Figure 3, could also indicate that the genetic units contributing to genetic similarity are not the same as the genetic units that contribute to similarity in cortical thickness (Boyle, Li, and Pritchard 2017). Some care should be taken with this interpretation, however, due to the highly complex time- and location (i.e. region)-specific influence of genetic factors on cortical thickness (Kang et al. 2011; Strike et al. 2019; van der Meer and Kaufmann 2022), which might not be captured well by a coarse similarity in cortical thickness across all vertices. The power to detect covariances in the present study might also be low due to the overall small estimates of morphometricity. In the original CORE GREML paper by Zhou et al. (2020), ten traits with high heritability were selected to maximize the power to detect genome-transcriptome covariance. A recent paper by Owens et al. (2021) showed that small effect sizes are generally expected in the ABCD study, which might make sound inference regarding gene-morphology covariance complicated in this sample.

### Limitations

The present paper has three limitations of particular importance. First, treating morphological and genetic effects as random avoids the issue of exhausting statistical power on hypothesis testing corrections for individual SNPs or vertices, but comes at the cost of spatial resolution, i.e. we cannot decipher which parts of the brain that contributed to variation in a trait.

Second, the present study is cross-sectional, representing only a snapshot of the child brain at a single point in time. It is possible that the link between individual similarity in cortical thickness and individual differences in traits is better understood looking at change over time (Foulkes and Blakemore 2018; Rakesh et al. 2023), or that the sensitivity of cortical thickness to relevant outcome variables increases as individuals age (Mewton et al. 2022). A promising avenue might involve the calculation of separate BRMs for multiple timepoints and look for changes in interindividual differences in cortical thickness or other morphological measures across time, and how these changes relate to both changes in the influence of genetic factors and in observable traits.

Third, any type of neuroimaging measure can be expressed as a relatedness matrix. In the present paper, we limited our approach to cortical thickness. To better capture the strength afforded by the multimodal approach of large-scale imaging studies, future studies should seek to integrate the information afforded by multiple imaging derived phenotypes.

### Concluding remarks

Here, we employed methods from statistical genetics to capture the association between cortical morphology and traits spanning the child phenome. Within the same linear mixed model framework, we assessed the effects of genetic similarity and its potential association with morphological similarity. Overall, associations with morphology were mostly limited to anthropometric traits, although some associations with socioeconomic status were also observed. The estimated contribution of genetic effects to trait variance was at the lower end of what is commonly found, possibly attributable to the age and ethnic makeup of the sample. No significant covariance between the effects of cortical morphology and genetic effects was found. Future studies should seek to better integrate information from different imaging derived measures beyond cortical thickness.

## Acknowledgements

This project was funded by research grants from the Research Council of Norway (Grant Nos. L.T.W: 249795, 300767. C.K.T: 288083, 323951. E.Y: 288083, 262177, 336078), the South-Eastern Norway Regional Health Authority (Grant Nos. L.T.W: 2014097, 2015073, 2016083, 2018076, 2019101. C.K.T: 2019069, 2021070, 2023012, 500189. D.A: 2019107, 2020086), the Norwegian ExtraFoundation for Health and Rehabilitation (L.T.W: Grant No. 2015/FO5146), KG Jebsen Stiftelsen, ERA-Net Cofund through the ERA PerMed project IMPLEMENT, the European Research Council under the European Union s Horizon 2020 research and Innovation program (L.T.W: ERC StG Grant No. 802998), and the European Research Council under the Horizon Europa program (E.Y: ERC CoG Grant No. 101045526). Views and opinions expressed are however those of the author(s) only and do not necessarily reflect those of the European Union or the European Research Executive Agency (REA). Neither the European Union nor the granting authority can be held responsible for them.

The work was performed on the Service for Sensitive Data (TSD) platform, owned by the University of Oslo, operated, and developed by the TSD service group at the University of Oslo IT-Department (USIT).

Data used in the preparation of this article were obtained from the Adolescent Brain Cognitive Development^SM^ (ABCD) Study (https://abcdstudy.org), held in the NIMH Data Archive (NDA). This is a multisite, longitudinal study designed to recruit more than 10,000 children aged 9-10 and follow them over 10 years into early adulthood. The ABCD Study® is supported by the National Institutes of Health and additional federal partners under award numbers U01DA041048, U01DA050989, U01DA051016, U01DA041022, U01DA051018, U01DA051037, U01DA050987, U01DA041174, U01DA041106, U01DA041117, U01DA041028, U01DA041134, U01DA050988, U01DA051039, U01DA041156, U01DA041025, U01DA041120, U01DA051038, U01DA041148, U01DA041093, U01DA041089, U24DA041123, U24DA041147. A full list of supporters is available at https://abcdstudy.org/federal-partners.html. A listing of participating sites and a complete listing of the study investigators can be found at https://abcdstudy.org/consortium_members/. ABCD consortium investigators designed and implemented the study and/or provided data but did not necessarily participate in the analysis or writing of this report. This manuscript reflects the views of the authors and may not reflect the opinions or views of the NIH or ABCD consortium investigators.

The ABCD data repository grows and changes over time. The ABCD data used in this report came from ABCD release 3.0 (NDA Study 901, DOI 10.15154/1519007).

## Notes

### Competing Interest Statement

The authors have declared no competing interest.

## References

Alnæs, Dag, Tobias Kaufmann, Andre F. Marquand, Stephen M. Smith, and Lars T. Westlye. 2020. “Patterns of Sociocognitive Stratification and Perinatal Risk in the Child Brain.” Proceedings of the National Academy of Sciences of the United States of America 117 (22): 12419–27.

Auchter, Allison M., Margie Hernandez Mejia, Charles J. Heyser, Paul D. Shilling, Terry L. Jernigan, Sandra A. Brown, Susan F. Tapert, and Gayathri J. Dowling. 2018. “A Description of the ABCD Organizational Structure and Communication Framework.” Developmental Cognitive Neuroscience 32 (August): 8–15.

Barnes, Josephine, Gerard R. Ridgway, Jonathan Bartlett, Susie M. D. Henley, Manja Lehmann, Nicola Hobbs, Matthew J. Clarkson, David G. MacManus, Sebastien Ourselin, and Nick C. Fox. 2010. “Head Size, Age and Gender Adjustment in MRI Studies: A Necessary Nuisance?” NeuroImage 53 (4): 1244–55.

Benjamini, Yoav, and Yosef Hochberg. 1995. “Controlling the False Discovery Rate: A Practical and Powerful Approach to Multiple Testing.” Journal of the Royal Statistical Society 57 (1): 289–300.

Bergen, Sarah E., Charles O. Gardner, and Kenneth S. Kendler. 2007. “Age-Related Changes in Heritability of Behavioral Phenotypes Over Adolescence and Young Adulthood: A Meta-Analysis.” Twin Research and Human Genetics: The Official Journal of the International Society for Twin Studies 10 (3): 423–33.

Botvinik-Nezer, Rotem, and Tor D. Wager. 2022. “Reproducibility in Neuroimaging Analysis: Challenges and Solutions.” Biological Psychiatry: Cognitive Neuroscience and Neuroimaging, December. https://doi.org/10.1016/j.bpsc.2022.12.006.

Boyle, Evan A., Yang I. Li, and Jonathan K. Pritchard. 2017. “An Expanded View of Complex Traits: From Polygenic to Omnigenic.” Cell 169 (7): 1177–86.

Casey, B. J., Tariq Cannonier, May I. Conley, Alexandra O. Cohen, Deanna M. Barch, Mary M. Heitzeg, Mary E. Soules, et al. 2018. “The Adolescent Brain Cognitive Development (ABCD) Study: Imaging Acquisition across 21 Sites.” Developmental Cognitive Neuroscience 32 (August): 43–54.

Cheesman, Rosa, Espen Moen Eilertsen, Yasmin I. Ahmadzadeh, Line C. Gjerde, Laurie J. Hannigan, Alexandra Havdahl, Alexander I. Young, et al. 2020. “How Important Are Parents in the Development of Child Anxiety and Depression? A Genomic Analysis of Parent-Offspring Trios in the Norwegian Mother Father and Child Cohort Study (MoBa).” BMC Medicine 18 (1): 284.

Conomos, Matthew P., Alexander P. Reiner, Bruce S. Weir, and Timothy A. Thornton. 2016. “Model-Free Estimation of Recent Genetic Relatedness.” American Journal of Human Genetics 98 (1): 127–48.

Couvy-Duchesne, Baptiste, Lachlan T. Strike, Futao Zhang, Yan Holtz, Zhili Zheng, Kathryn E. Kemper, Loic Yengo, et al. 2020. “A Unified Framework for Association and Prediction from Vertex-Wise Grey-Matter Structure.” Human Brain Mapping 41 (14): 4062–76.

Dennis, Evan, Peter Manza, and Nora D. Volkow. 2022. “Socioeconomic Status, BMI, and Brain Development in Children.” Translational Psychiatry 12 (1): 33.

Donati, Georgina, Iroise Dumontheil, Oliver Pain, Kathryn Asbury, and Emma L. Meaburn. 2021. “Evidence for Specificity of Polygenic Contributions to Attainment in English, Maths and Science during Adolescence.” Scientific Reports 11 (1): 3851.

Eilertsen, Espen M., Rosa Cheesman, Ziada Ayorech, Espen Røysamb, Jean-Baptiste Pingault, Pål R. Njølstad, Ole A. Andreassen, et al. 2022. “On the Importance of Parenting in Externalizing Disorders: An Evaluation of Indirect Genetic Effects in Families.” Journal of Child Psychology and Psychiatry, and Allied Disciplines 63 (10): 1186–95.

Estrada, Eduardo, Emilio Ferrer, Francisco J. Román, Sherif Karama, and Roberto Colom. 2019. “Time-Lagged Associations between Cognitive and Cortical Development from Childhood to Early Adulthood.” Developmental Psychology 55 (6): 1338–52.

Fernandez-Cabello, Sara, Dag Alnæs, Dennis van der Meer, Andreas Dahl, Madelene Holm, Rikka Kjelkenes, Ivan I. Maximov, et al. 2022. “Associations between Brain Imaging and Polygenic Scores of Mental Health and Educational Attainment in Children Aged 9-11.” NeuroImage 263 (September): 119611.

Fortin, Jean-Philippe, Nicholas Cullen, Yvette I. Sheline, Warren D. Taylor, Irem Aselcioglu, Philip A. Cook, Phil Adams, et al. 2018. “Harmonization of Cortical Thickness Measurements across Scanners and Sites.” NeuroImage 167 (February): 104–20.

Foulkes, Lucy, and Sarah-Jayne Blakemore. 2018. “Studying Individual Differences in Human Adolescent Brain Development.” Nature Neuroscience 21 (3): 315–23.

Frangou, Sophia, Amirhossein Modabbernia, Steven C. R. Williams, Efstathios Papachristou, Gaelle E. Doucet, Ingrid Agartz, Moji Aghajani, et al. 2022. “Cortical Thickness across the Lifespan: Data from 17,075 Healthy Individuals Aged 3-90 Years.” Human Brain Mapping 43 (1): 431–51.

Fuhrmann, D., K. S. Madsen, L. B. Johansen, W. F. C. Baaré, and R. A. Kievit. 2022. “The Midpoint of Cortical Thinning between Late Childhood and Early Adulthood Differs between Individuals and Brain Regions: Evidence from Longitudinal Modelling in a 12-Wave Neuroimaging Sample.” NeuroImage 261 (November): 119507.

Fürtjes, Anna E., James H. Cole, Baptiste Couvy-Duchesne, and Stuart J. Ritchie. 2023. “A Quantified Comparison of Cortical Atlases on the Basis of Trait Morphometricity.” Cortex; a Journal Devoted to the Study of the Nervous System and Behavior 158 (January): 110–26.

Genon, Sarah, Simon B. Eickhoff, and Shahrzad Kharabian. 2022. “Linking Interindividual Variability in Brain Structure to Behaviour.” Nature Reviews. Neuroscience 23 (5): 307–18.

Giedd, Jay N., and Judith L. Rapoport. 2010. “Structural MRI of Pediatric Brain Development: What Have We Learned and Where Are We Going?” Neuron 67 (5): 728–34.

Gilmore, John H., Benjamin Langworthy, Jessica B. Girault, Jason Fine, Shaili C. Jha, Sun Hyung Kim, Emil Cornea, and Martin Styner. 2020. “Individual Variation of Human Cortical Structure Is Established in the First Year of Life.” Biological Psychiatry. Cognitive Neuroscience and Neuroimaging 5 (10): 971–80.

Hettwer, M. D., S. Larivière, B. Y. Park, O. A. van den Heuvel, L. Schmaal, O. A. Andreassen, C. R. K. Ching, et al. 2022. “Coordinated Cortical Thickness Alterations across Six Neurodevelopmental and Psychiatric Disorders.” Nature Communications 13 (1): 6851.

Jami, Eshim S., Anke R. Hammerschlag, Hill F. Ip, Andrea G. Allegrini, Beben Benyamin, Richard Border, Elizabeth W. Diemer, et al. 2022. “Genome-Wide Association Meta-Analysis of Childhood and Adolescent Internalizing Symptoms.” Journal of the American Academy of Child and Adolescent Psychiatry 61 (7): 934–45.

Jelenkovic, Aline, Reijo Sund, Yoon-Mi Hur, Yoshie Yokoyama, Jacob V. B. Hjelmborg, Sören Möller, Chika Honda, et al. 2016. “Genetic and Environmental Influences on Height from Infancy to Early Adulthood: An Individual-Based Pooled Analysis of 45 Twin Cohorts.” Scientific Reports 6 (June): 28496.

Kang, Hyo Jung, Yuka Imamura Kawasawa, Feng Cheng, Ying Zhu, Xuming Xu, Mingfeng Li, André M. M. Sousa, et al. 2011. “Spatio-Temporal Transcriptome of the Human Brain.” Nature 478 (7370): 483–89.

Knickmeyer, Rebecca C., Kai Xia, Zhaohua Lu, Mihye Ahn, Shaili C. Jha, Fei Zou, Hongtu Zhu, Martin Styner, and John H. Gilmore. 2017. “Impact of Demographic and Obstetric Factors on Infant Brain Volumes: A Population Neuroscience Study.” Cerebral Cortex 27 (12): 5616–25.

Lam, Max, Swapnil Awasthi, Hunna J. Watson, Jackie Goldstein, Georgia Panagiotaropoulou, Vassily Trubetskoy, Robert Karlsson, et al. 2020. “RICOPILI: Rapid Imputation for COnsortias PIpeLIne.” Bioinformatics 36 (3): 930–33.

Lashley, K. S. 1947. “Structural Variation in the Nervous System in Relation to Behavior.” Psychological Review 54 (6): 325–34.

Lee, S. H., and J. H. J. van der Werf. 2016. “MTG2: An Efficient Algorithm for Multivariate Linear Mixed Model Analysis Based on Genomic Information.” Bioinformatics 32 (9): 1420–22.

Leys, Christophe, Christophe Ley, Olivier Klein, Philippe Bernard, and Laurent Licata. 2013. “Detecting Outliers: Do Not Use Standard Deviation around the Mean, Use Absolute Deviation around the Median.” Journal of Experimental Social Psychology 49 (4): 764–66.

Li, Yun R., and Brendan J. Keating. 2014. “Trans-Ethnic Genome-Wide Association Studies: Advantages and Challenges of Mapping in Diverse Populations.” Genome Medicine 6 (10): 91.

Marek, Scott, Brenden Tervo-Clemmens, Finnegan J. Calabro, David F. Montez, Benjamin P. Kay, Alexander S. Hatoum, Meghan Rose Donohue, et al. 2022. “Reproducible Brain-Wide Association Studies Require Thousands of Individuals.” Nature 603 (7902): 654–60.

Meer, Dennis van der, Oleksandr Frei, Tobias Kaufmann, Chi-Hua Chen, Wesley K. Thompson, Kevin S. O’Connell, Jennifer Monereo Sánchez, et al. 2020. “Quantifying the Polygenic Architecture of the Human Cerebral Cortex: Extensive Genetic Overlap between Cortical Thickness and Surface Area.” Cerebral Cortex 30 (10): 5597–5603.

Meer, Dennis van der, and Tobias Kaufmann. 2022. “Mapping the Genetic Architecture of Cortical Morphology through Neuroimaging: Progress and Perspectives.” Translational Psychiatry 12 (1): 447.

Mewton, Louise, Briana Lees, Lindsay M. Squeglia, Miriam K. Forbes, Matthew Sunderland, Robert Krueger, Forrest C. Koch, et al. 2022. “The Relationship between Brain Structure and General Psychopathology in Preadolescents.” Journal of Child Psychology and Psychiatry, and Allied Disciplines 63 (7): 734–44.

Mills, Kathryn L., Kimberly D. Siegmund, Christian K. Tamnes, Lia Ferschmann, Lara M. Wierenga, Marieke G. N. Bos, Beatriz Luna, Chun Li, and Megan M. Herting. 2021. “Inter-Individual Variability in Structural Brain Development from Late Childhood to Young Adulthood.” NeuroImage 242 (November): 118450.

Momin, Md Moksedul, Jisu Shin, Soohyun Lee, Buu Truong, Beben Benyamin, and S. Hong Lee. 2023. “A Method for an Unbiased Estimate of Cross-Ancestry Genetic Correlation Using Individual-Level Data.” Nature Communications 14 (1): 1–13.

Moore, K. A., D. E. Myers, D. R. Morrison, C. W. Nord, B. Brown, and B. Edmonston. 1993. “Age at First Childbirth and Later Poverty.” Journal of Research on Adolescence: The Official Journal of the Society for Research on Adolescence 3 (4): 393–422.

Oehlert, Gary W. 1992. “A Note on the Delta Method.” The American Statistician 46 (1): 27–29.

Owens, Max M., Alexandra Potter, Courtland S. Hyatt, Matthew Albaugh, Wesley K. Thompson, Terry Jernigan, Dekang Yuan, Sage Hahn, Nicholas Allgaier, and Hugh Garavan. 2021. “Recalibrating Expectations about Effect Size: A Multi-Method Survey of Effect Sizes in the ABCD Study.” PloS One 16 (9): e0257535.

Rakesh, Divyangana, Sarah Whittle, Margaret A. Sheridan, and Katie A. McLaughlin. 2023. “Childhood Socioeconomic Status and the Pace of Structural Neurodevelopment: Accelerated, Delayed, or Simply Different?” Trends in Cognitive Sciences, May. https://doi.org/10.1016/j.tics.2023.03.011.

Restrepo-Méndez, María Clara, Debbie A. Lawlor, Bernardo L. Horta, Alicia Matijasevich, Iná S. Santos, Ana M. B. Menezes, Fernando C. Barros, and Cesar G. Victora. 2015. “The Association of Maternal Age with Birthweight and Gestational Age: A Cross-Cohort Comparison.” Paediatric and Perinatal Epidemiology 29 (1): 31–40.

Sabuncu, Mert R., Tian Ge, Avram J. Holmes, Jordan W. Smoller, Randy L. Buckner, Bruce Fischl, and Alzheimer’s Disease Neuroimaging Initiative. 2016. “Morphometricity as a Measure of the Neuroanatomical Signature of a Trait.” Proceedings of the National Academy of Sciences of the United States of America 113 (39): E5749–56.

Schmitt, J. Eric, Michael C. Neale, Bilqis Fassassi, Javier Perez, Rhoshel K. Lenroot, Elizabeth M. Wells, and Jay N. Giedd. 2014. “The Dynamic Role of Genetics on Cortical Patterning during Childhood and Adolescence.” Proceedings of the National Academy of Sciences of the United States of America 111 (18): 6774–79.

Shadrin, Alexey A., Tobias Kaufmann, Dennis van der Meer, Clare E. Palmer, Carolina Makowski, Robert Loughnan, Terry L. Jernigan, et al. 2021. “Vertex-Wise Multivariate Genome-Wide Association Study Identifies 780 Unique Genetic Loci Associated with Cortical Morphology.” NeuroImage 244 (December): 118603.

Strike, Lachlan T., Narelle K. Hansell, Baptiste Couvy-Duchesne, Paul M. Thompson, Greig I. de Zubicaray, Katie L. McMahon, and Margaret J. Wright. 2019. “Genetic Complexity of Cortical Structure: Differences in Genetic and Environmental Factors Influencing Cortical Surface Area and Thickness.” Cerebral Cortex 29 (3): 952–62.

Tam, Vivian, Nikunj Patel, Michelle Turcotte, Yohan Bossé, Guillaume Paré, and David Meyre. 2019. “Benefits and Limitations of Genome-Wide Association Studies.” Nature Reviews. Genetics 20 (8): 467–84.

Tomasi, Dardo, and Nora D. Volkow. 2021. “Associations of Family Income with Cognition and Brain Structure in USA Children: Prevention Implications.” Molecular Psychiatry 26 (11): 6619–29.

Trzaskowski, M., J. Yang, P. M. Visscher, and R. Plomin. 2014. “DNA Evidence for Strong Genetic Stability and Increasing Heritability of Intelligence from Age 7 to 12.” Molecular Psychiatry 19 (3): 380–84.

Volkow, Nora D., George F. Koob, Robert T. Croyle, Diana W. Bianchi, Joshua A. Gordon, Walter J. Koroshetz, Eliseo J. Pérez-Stable, et al. 2018. “The Conception of the ABCD Study: From Substance Use to a Broad NIH Collaboration.” Developmental Cognitive Neuroscience 32 (August): 4–7.

Walhovd, Kristine B., Stine Kleppe Krogsrud, Inge K. Amlien, Øystein Sørensen, Yunpeng Wang, Anne Cecilie Sjøli Bråthen, Knut Overbye, et al. 2023. “Back to the Future: Omnipresence of Fetal Influence on the Human Brain through the Lifespan.” bioRxiv. https://doi.org/10.1101/2022.12.02.514196.

Westlin, Christiana, Jordan E. Theriault, Yuta Katsumi, Alfonso Nieto-Castanon, Aaron Kucyi, Sebastian F. Ruf, Sarah M. Brown, et al. 2023. “Improving the Study of Brain-Behavior Relationships by Revisiting Basic Assumptions.” Trends in Cognitive Sciences 27 (3): 246–57.

Yang, Jian, Beben Benyamin, Brian P. McEvoy, Scott Gordon, Anjali K. Henders, Dale R. Nyholt, Pamela A. Madden, et al. 2010. “Common SNPs Explain a Large Proportion of the Heritability for Human Height.” Nature Genetics 42 (7): 565–69.

Yang, Jian, S. Hong Lee, Michael E. Goddard, and Peter M. Visscher. 2011. “GCTA: A Tool for Genome-Wide Complex Trait Analysis.” American Journal of Human Genetics 88 (1): 76–82.

Yengo, Loïc, Sailaja Vedantam, Eirini Marouli, Julia Sidorenko, Eric Bartell, Saori Sakaue, Marielisa Graff, et al. 2022. “A Saturated Map of Common Genetic Variants Associated with Human Height.” Nature 610 (7933): 704–12.

Yu, Jianming, Gael Pressoir, William H. Briggs, Irie Vroh Bi, Masanori Yamasaki, John F. Doebley, Michael D. McMullen, et al. 2006. “A Unified Mixed-Model Method for Association Mapping That Accounts for Multiple Levels of Relatedness.” Nature Genetics 38 (2): 203–8.

Yu, Zhaoxia, Michele Guindani, Steven F. Grieco, Lujia Chen, Todd C. Holmes, and Xiangmin Xu. 2022. “Beyond T Test and ANOVA: Applications of Mixed-Effects Models for More Rigorous Statistical Analysis in Neuroscience Research.” Neuron 110 (1): 21–35.

Zhou, Xuan, Hae Kyung Im, and S. Hong Lee. 2020. “CORE GREML for Estimating Covariance between Random Effects in Linear Mixed Models for Complex Trait Analyses.” Nature Communications 11 (1): 4208.

