## Supplementary material for "Genetic and brain similarity independently predict childhood anthropometrics and socioeconomic markers"

**Supplementary figure 1.** Data processing flow

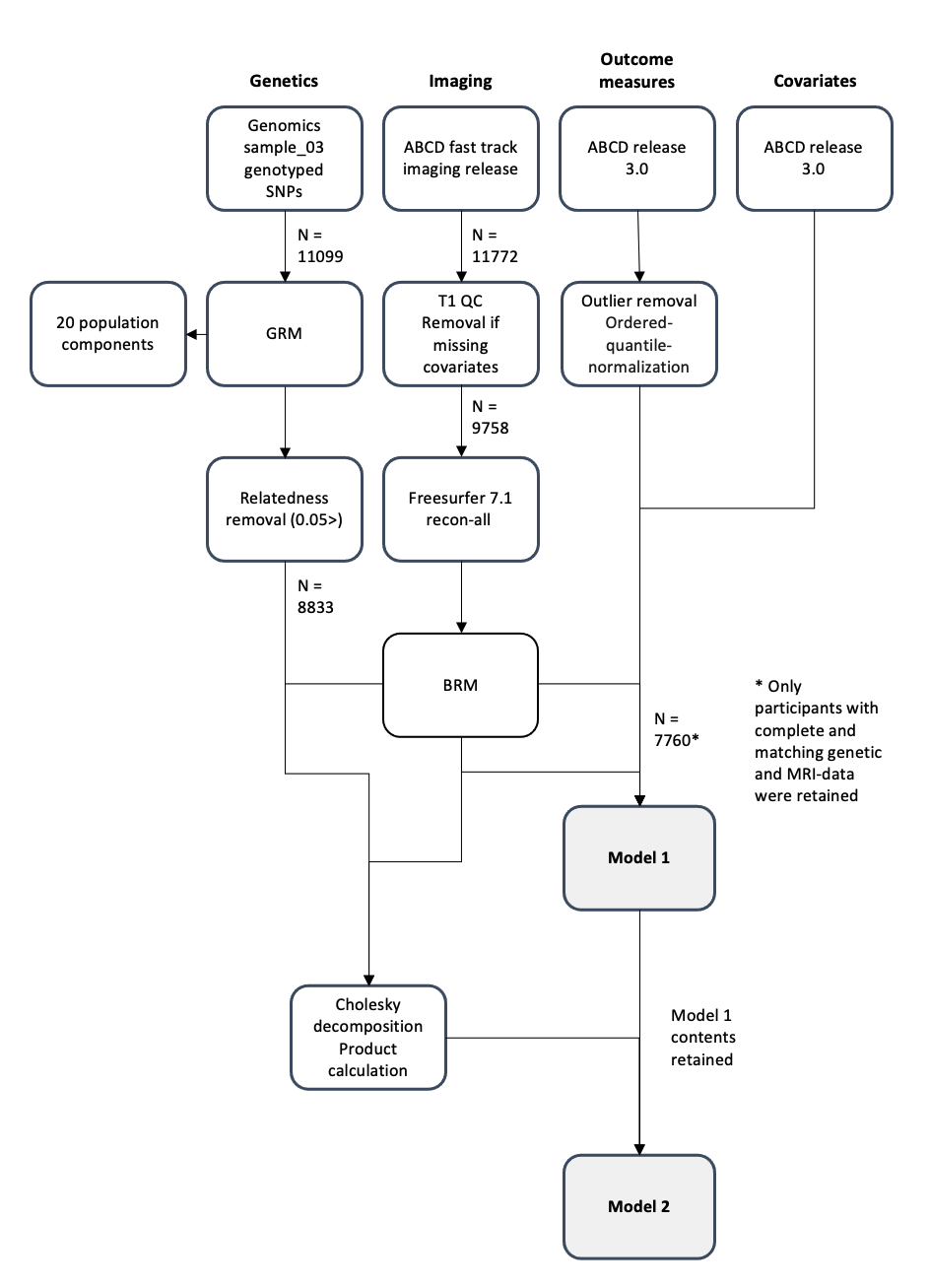

**Supplementary figure 2**. Mean cortical thickness at each scanner before (A) and after (B) the neuroCombat harmonization procedure.

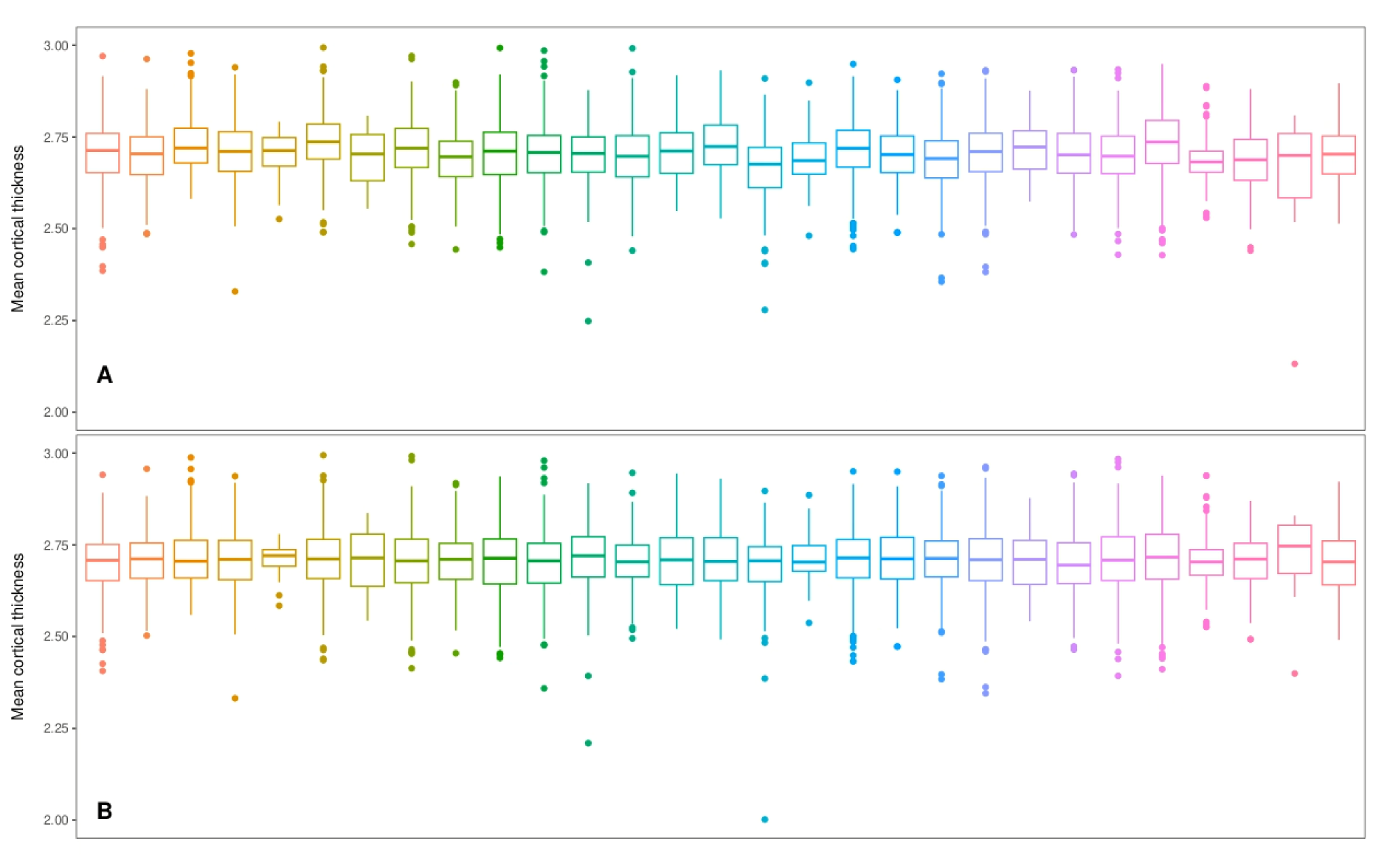

**Supplementary table 1**. All Model 1 outcomes.

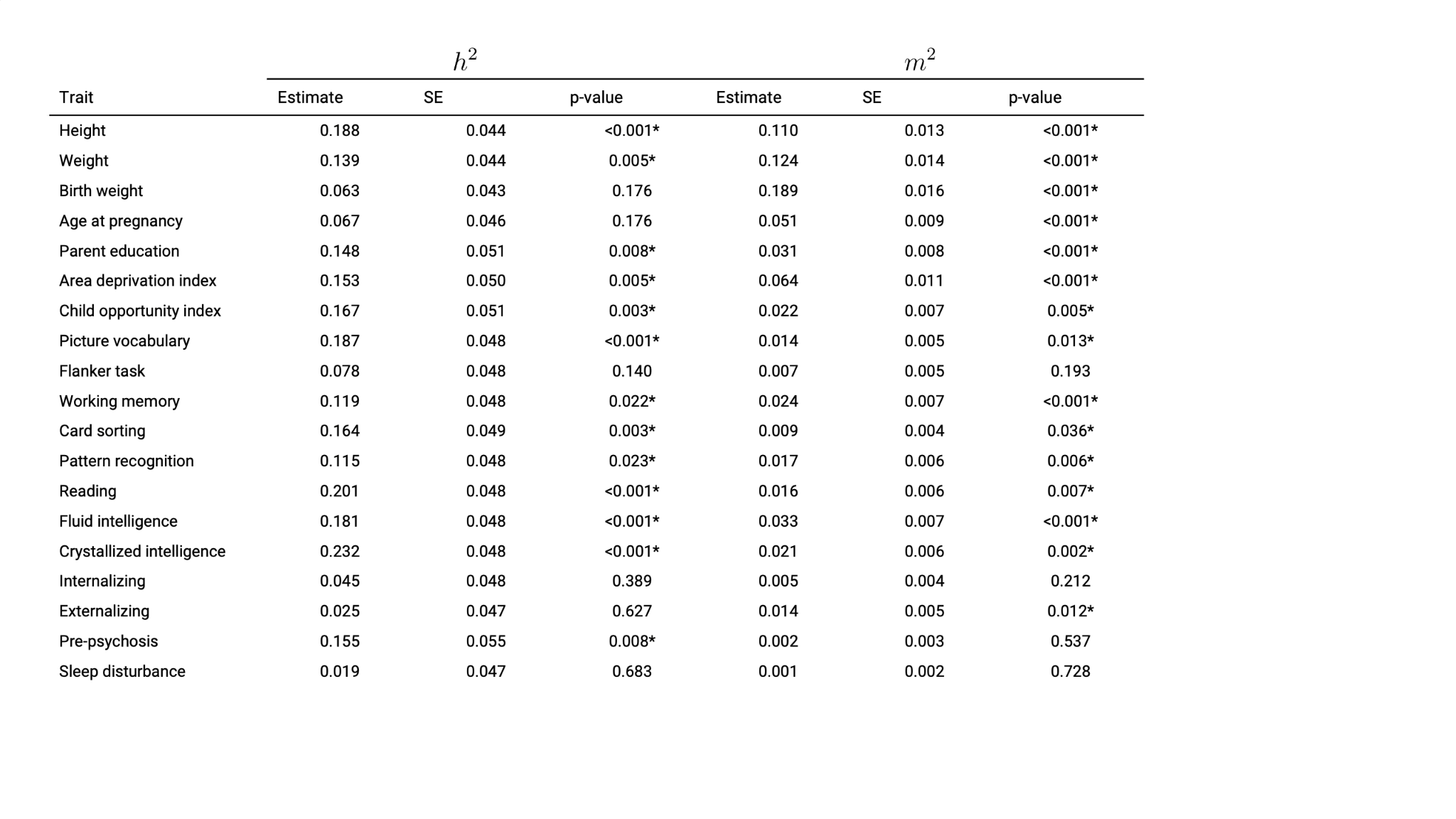
